# First detection of *Leptospira santarosai* in the reproductive track of a boar: a potential threat to swine production and public health

**DOI:** 10.1101/2022.04.20.488867

**Authors:** Eduardo A. Diaz, Ligia Luna, Ana Burgos-Mayorga, Gustavo Donoso, Diego A. Guzman, María Ines Baquero, Talima Pearson, Veronica Barragan

## Abstract

**Background:** Leptospirosis causes significant economic losses and is an occupational risk in the swine industry, especially in developing tropical regions where social and geoclimatic conditions are favorable for the transmission of this disease. Although vaccination can reduce infection risk, efficacy is diminished if local genetic and antigenic variants of the pathogen are not accounted for in the vaccine. Identifying and characterizing strains that circulate in different populations is therefore critical for public health mitigation practices.

**Methodology/Principal findings:** Our study was conducted on a rural breeding farm in Ecuador, where we identified, for the first time, *Leptospira santarosai* in the kidneys, testicles, and ejaculate of a vaccinated boar. *L. santarosai* was detected with a PCR assay that targets *lipL32*, and identified by target MLST gene sequencing using an Oxford Nanopore MinION sequencer.

**Conclusions/Significance:** As *L. santarosai* is pathogenic in other livestock species and humans, our finding highlights the need to evaluate the prevalence and epidemiological significance of this pathogen in pigs. In addition, further studies are needed to identify and characterize local serovars that may impact diagnosis and vaccination programs to better control leptospirosis in pigs and spillover into the human population.

**Author summary:** Leptospirosis poses a significant threat to human and animal health. In tropical countries, leptospirosis is very common, and responsible of economic losses in the livestock industry. In peridomestic and rural farms, the spillover of leptospira to humans is particularly likely as humans live and work in close proximity to animals. Although animal vaccination can reduce risk of infection, efficacy is diminished when local variants are not included in the vaccine. This report describes, for the first time, the presence of *Leptospira santarosai* in the reproductive tract of a vaccinated domestic boar from a rural farm in Ecuador. We detected the pathogen in its semen and urine, and despite no tissue damage, was observed in the kidneys, testes or epididymis. The farm veterinarian reported reproductive problems in sows inseminated with the semen from this boar. Our results highlight the importance of recognizing locally circulating serovars and species so that they can be included in vaccines to prevent infection and disease. Effective control of leptospirosis in livestock not only reduces economic losses for breeders, but also reduces the risk of infection and disease in humans.

## Introduction

Leptospirosis is a reemerging zoonosis with worldwide distribution and a significant impact on livestock production. The disease is frequently associated with reproductive disorders, including embryonic resorption, fetal mummification, stillbirths, or neonatal mortality causing significant economic losses. Furthermore, leptospirosis in farmers, slaughterhouse workers, and veterinarians is very common and occurs through direct contact with urine or tissues from infected animals, or indirectly through contaminated soil and water [1]. Unfortunately, knowledge about the disease in livestock is biased towards intensive breeding industries in developed regions of the world, while the epidemiological characteristics of leptospirosis in developing nations remain unclear [2].

In most low-income tropical countries, leptospirosis is endemic and common. Ownership of a small number of peridomestic livestock is common in poor rural areas, and pigs are often part of this community. In these situations, pigs are often raised under poor sanitary conditions without veterinary guidance [3,4]. Importantly, the social and geoclimatic characteristics of these areas are conducive to the transmission and maintenance of *Leptospira*. Indeed, in endemic regions, leptospira can persist in the urogenital tract of asymptomatic animals that excrete the bacteria in urine and genital fluids, providing a source of infection for susceptible hosts [5]. In these settings, spillover to humans is likely as humans and animals live in close proximity [2]. Although vaccination can reduce leptospira transmission, efficacy is commonly reduced due to low immunity against strains that are not represented in vaccines [6]. This is especially problematic in countries where very little information on circulating serovars is available and when local isolates are not available for inclusion in MAT tests for disease diagnosis. Given the high, uncharacterized diversity of pathogenic *Leptospira* in tropical developing countries, understanding the local epidemiology of leptospirosis is paramount for disease mitigation.

Recent studies in rural communities on the coast of Ecuador show that exposure to livestock is common and pigs may serve as an under-recognized source of high Leptospira diversity in the region [7–9]. Swine leptospirosis has historically been linked to exposure to urine from carrier animals, but there have been no links to the reproductive system in disease transmission [7,8,10,11].

Here, we present the case of a domestic boar raised in a rural area of Ecuador, that was found to excrete Leptospira in its semen. The boar had been vaccinated and showed no clinical signs of disease, but laboratory analysis identified a pathogenic Leptospira species that had not been previously described in the reproductive tract of pigs. We present serologic, molecular, and histopathologic data from this rare case of porcine genital carriage of pathogenic Leptospira. Our results reaffirm the need for a thorough understanding of the epidemiology of leptospirosis in endemic regions at the local level in order to implement appropriate preventive vaccination and improve disease control programs [12].

## Materials and methods

### Boar information

The subject under study was a 2-year-old Landrace/Yorkshire crossbreed boar in its reproductive stage, maintained in a rural pig-breeding farm located in the suburbs of Quito. Within the farm, animals are kept in individual pens, have routine veterinary visits, a balanced diet, free access to feed and water, and occasional plague management and cleaning. Animals on the farm are vaccinated every six months, receiving an anti-leptospiral vaccine against *Leptospira interrogans* serovars Bratislava, Canicola, Grippotyphosa, Hardjo, Icterohaemorrhagie, and Pomona (Farrow sure®, Zoetis). The last leptospirosis vaccination for the boar occurred 1 month prior to the collection of the first diagnostic blood sample (June 2019).

The boar’s semen was collected weekly and provided to external producers, and used for artificial insemination within the farm. In February 2019, two sows inseminated with semen from this boar had reproductive problems (reabsorption and repetition of estrus). Later, a second insemination of the same sows resulted in mummification during farrowing. In addition, external pork producers who bought the semen, also reported fetus mummifications. These reports, coupled with blood observed in the semen, came to the attention of the farm veterinarian who collected diagnostic specimens to test for leptospirosis. Two additional indicators for leptospirosis and possible circulation of pathogenic Leptospira were observed by the farm veterinarian: 1. rat droppings were frequently found inside and outside the pens, and 2. one year earlier (2018) a different farrowed sow showed blood in her urine. This sow was impregnated with a different boar. The serum sample from this animal showed high Microscopic Agglutination Test (MAT) antibodies titers, however the offspring were negative for the MAT. We were unable to access the samples or results from this sow.

### Sample collection

Blood samples were collected from the jugular vein of the boar on two different dates (Sample S1 on July 2019, and Sample S2 on August 2019), and sent to the National Reference Laboratory for Animal Diagnostics (AGROCALIDAD) to be tested with a Microscopic Agglutination Test (MAT). Considering that *Leptospira* in urine can be intermittently excreted [13], four urine samples were collected by spontaneous micturition on different dates: September 9^th^ and 26^th^, October 1^st^ 2019, and February 20^th^ 2020. Three semen samples were collected on September 20^th^, 26^th^ and October 1^st^ 2019. Each semen sample was divided into spermatic (semen) and post-spermatic (dense discharge from the Cowper glands and prostate) fractions and placed into sterile tubes.

On February 20^th^ 2020, the farm owners culled the boar, allowing us to collect tissue samples. Kidney, testicle, and epididymis samples were placed in sterile tubes with 80% ethanol for molecular detection of *Leptospira*, and in 10% buffered formalin for histopathological examination. Samples were transported on ice to the microbiology laboratory of the Universidad San Francisco de Quito and kept at −20°C before analyses.

### *Leptospira* detection

DNA from tissue and fluids was extracted using the DNeasy Blood and Tissue kit - Qiagen, CA, USA for semen and urine, and Purelink Genomic DNA kit - Invitrogen, Carlsbad, CA, USA for kidney and testicle tissue. Different kits were used because of inconsistent availability in Ecuador. Detection of the *lipL32* gene was used to define Leptospira DNA positivity [14]. Positive samples were subjected to a second round of PCR using MLST primers [15]. MLST amplicons were sequenced using a portable Oxford Nanopore MinION sequencer. Library preparation was performed using the Barcoding kit (SQK-RBK004 - Oxford Nanopore Technologies), and loaded into a MinION Flowcell (FLO-MIN 106). Guppy (version 3.4.5) was used for basecalling of FAST5 files. Porechop (version 0.2.4) (https://github.com/rrwick/Porechop) was used to perform demultiplexing and adapter removal, and Nanoplot was used to determine sequence quality (http://nanoplot.bioinf.be/). Leptospira amplicons were filtered using the BLAST command line tool [16], aligned using minimap2 [17], and visualized using Tablet [18]. Consensus sequences were obtained using online EMBL-EBI search and sequence analysis tools (https://www.ebi.ac.uk/Tools/msa/emboss_cons/), and identification of *Leptospira* species was confirmed using the online BLAST tool (https://blast.ncbi.nlm.nih.gov/Blast.cgi).

### Histopathological analysis

Samples submitted for histopathological analysis were embedded in paraffin [19]. Blocks were cut to 4 μm, stained with hematoxylin-eosin, and observed under light microscopy [19].

### Ethics statement

Verbal consent from the owner was provided throughout the entire study. Samples were collected under the permit issued by the Animal Bioethics Committee at Universidad San Francisco de Quito (Official Letter 2019-004-a), and molecular detection of *Leptospira* was performed under the permit: Contrato Marco de Acceso a los Recursos Genéticos (MAE-DNB-CM-2018-0106).

## Results

MAT on serum samples gave positive results with titters of 1:100 for serovar Canicola in sample S1, and 1:100 for serovars Pomona and Hardjo in sample S2 (Table S1).

Leptospira DNA was detected in semen (spermatic and post-spermatic fraction), and kidney and testicle tissues (Table 1). We were able to sequence four MLST genes (*icdA, lipL41, secY*, and *16S rDNA*) from a semen sample. These sequences were indicative of *Leptospira santarosai* with 99% identity (Data available at https://www.ncbi.nlm.nih.gov Bioproject ID: PRJNA741491, Biosample ID: SAMN1989635).

**Table 1.** Positivity of pathogenic *Leptospira* (amplification of *lipL32* gene) in urine, semen, and tissue samples.

| *lipL32* qPCR Results |  |  |  |  |  |  |
| --- | --- | --- | --- | --- | --- | --- |
| Date | Urine | Spermatic Fraction | Post spermatic fraction | Kidney | Epididymis | Testicles |
| 20/09/2019 | Negative | Negative | Positive<br>(Ct=32,15) | — | — | — |
| 26/09/2019 | Negative | Positive<br>(Ct=15,95) | Positive<br>(Ct=21,21) | — | — | — |
| 01/10/2019 | Negative | Negative | Positive<br>(Ct=32,57) | — | — | — |
| 20/02/2020 | Negative | — | — | Positive<br>(Ct=32,38) | Negative | Positive<br>(Ct=26,87) |
| % Positivity | 0 | 33 | 100 | 100 | 0 | 100 |

Interestingly, histopathology from kidney, testicle, and epididymis stained with hematoxylin-eosin, showed intact tissue with no microscopic lesions (Fig 1). Also, no gross lesions were observed in the organs during necropsy.

**Fig 1.**
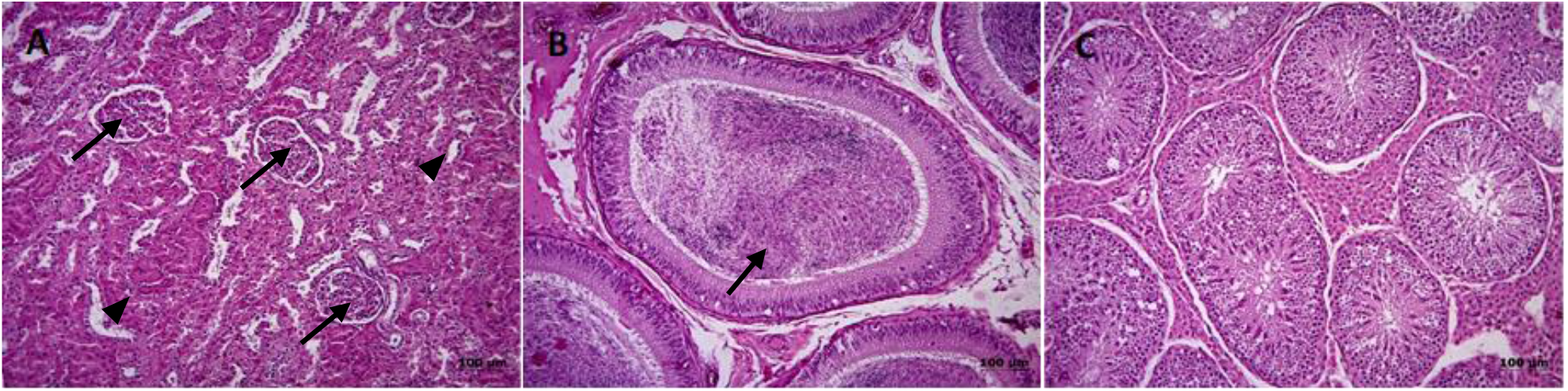
Histopathology of the boar organs. (A) Histology of the kidney presenting normal glomeruli (arrows) and tubules (arrowheads), showing no sign of inflammation. (B) Normal histology of the epididymis, exhibiting abundant spermatozoa in the lumen (arrow). (C) Normal histology of the testicle, physiological development of cell populations and normal architecture are maintained. No signs of inflammation are evident.

## Discussion

Genital leptospirosis can go unnoticed, compromising the reproductive productivity of herds over long periods of time [20]. However, the severity of the disease varies depending on the infecting strain and the affected species [2]. This research is the first record of *L. santarosai* in the reproductive system of a boar; specifically in testes and semen. We were also able to detect the bacteria in kidney tissue, but not in urine, probably because the animal was not excreting detectable amounts of leptospires at the time of urine collection and the focus of infection seems to have been in the reproductive tract. *Leptospira santarosai* have been reported in pig urine samples from Ecuador [7]. Our current findings are of particular interest because they indicate that reproductive and urinary aspects of the urogenital track of boars can both be colonized with *L. santarosai*, and present two routes of shedding and transmission.

Our research was carried out after observing that some sows, inseminated with semen from the same donor, had reproductive failures. The donor boar did not show clinical signs of leptospirosis, and the histopathological samples did not show evidence of tissue damage in the testes or epididymis. However, PCR and sequencing identified *L. santarosai* in testicles and semen samples. As previously reported for bulls [21], rams [22] or stallions [23] without apparent clinical signs, the presence of pathogenic *Leptospira* species DNA in semen suggests the potential venereal transmission of this pathogen. *Leptospira interrogans* and *L. kirschneri* have also been isolated from the reproductive tract of boars in Italy [24]. In the present case, the history of failed pregnancies provide evidence that leptospires were transmitted from the boar to sows via semen during artificial insemination (AI). In fact, *L. santarosai* has previously been associated with AI-transmitted bovine genital leptospirosis in other Latin American countries [20]. AI is a useful tool to introduce superior genes into herds and reduces the risk of injury and disease transmission through natural mating. However, semen can be contaminated with pathogens like *Leptospira* spp. [25]. The best strategy to prevent diseases transmitted by AI is to use pathogen-free boars, regularly monitoring animals and semen, and maintaining biosecurity strategies such as rodent control [26]. Indeed, the risk of *Leptospira* occurrence in the semen of boars from large commercial farms is low [27]. However, as sanitary conditions can affect AI, this method is a risk factor associated with leptospirosis on pig farms [28,29]. To our knowledge, prior to our study, the boar never tested positive for leptospirosis, however the presence of rodents in and around the farm put the animal at risk of infection.

Vaccination is one of the main strategies used to limit the spread of leptospires in herds. Currently, polyvalent vaccines include the most frequent serovars. However, vaccines are less effective in sites where local serovars are not non-included in the vaccine [30]. This is consistent with the fact that the boar under study was infected despite being vaccinated. Specifically, the vaccine used on this boar includes six serovars (Bratislava, Canicola, Grippotyphosa, Hardjo, Icterohaemorrhagiae and Pomona), while 15 different serovars have been identified in *L. santarosai* (Alice, Atlantae, Babudieri, Bananal, Batavidae, Beye, Canalzonae, Georgia, Guaricura, Kremastos, Peru, Pyrogenes, Shermani, Szwajizak and Tabaquite) but were not contained in the vaccine used on the farm [31–33].

Identification and removal of infected animals is used to control many infectious diseases, but is of limited value when the subclinical form is the most common presentation, and tests do not reliably identify carrier animals [2]. The MAT is the most common serological screening method used for leptospirosis. It is performed by incubating patient serum with various serovars of leptospires with any reacting serovar being indicative of the infecting serovar. Confirmation of an active infection is made by testing a second sample and demonstrating an increase in antibody titer. The MAT has the advantage of being serovar specific, but is prone to false negative results if the panel does not contain representative antigens of local serovars [6]. Furthermore, the MAT results must be interpreted with caution as it cannot discriminate between antibodies resulting from infection or vaccination, and high titers are not necessarily indicative of infection [34]. In our case, the boar MAT results show titers of 1:100 against three different serovars (Canicola, Pomona, and Hardjo), but this is undoubtedly due to vaccine-based immunity [6]. It is impossible to know if the animal would have shown any response to a serovar of *L. santarosai* because the reference diagnostic laboratory, where the MAT was performed, does not use *L. santarosai* serovars or any characterized local isolates. Molecular methods are also important tools for diagnosis in animals that do not show serological responses [35]. As previously reported for other domestic species [21–23], our results show that PCR and sequencing are important tools for the detection and characterization of leptospires in semen at swine artificial insemination centers. However, PCR and amplicon sequencing typically provides only species level identification and does not allow for serovar identification, limiting translational utility. This information is crucial for a better understanding of the epidemiology, the utility of diagnostic tests, and the development of new vaccines [36]. Based on these findings, we consider that the combined use of MAT as a screening test, followed by PCR and amplicon sequencing for the direct detection of, and characterization of *Leptospira* spp. was adequate for the identification of carrier animals, but bacteriological isolation of local serovars is critical for increasing the accuracy of MATs and improvement to vaccination strategies.

Leptospirosis is considered an underreported occupational disease, especially in developing countries [37]. Transmission among animals and humans through direct contact or indirectly through contaminated environments in low-tech peridomestic pig farms may be relatively common [38,39]. *Leptospira santarosai* have been previously identified in cases of human leptospirosis [33,40,41], and our findings reflect the potential risk of pigs on rural farms in low-income countries as a possible source of human and animal leptospirosis. Prevention and control measures for leptospirosis must be approached from a one-health perspective, however, a major limiting factor has been the lack of communication and cooperation between the human and animal healthcare communities [42]. This is the case of Ecuador, where leptospirosis is a notifiable human disease, but not a notifiable animal disease. In fact, the Ministry of Public Health reported 643 cases of human leptospirosis between 2016 and 2021, but there are no official reports of leptospirosis in cattle [43]. This information gap contributes to the lack of knowledge of the epidemiology of leptospirosis in the region.

## Conclusion

This is the first report on the detection and identification of a pathogenic *Leptospira* from the reproductive system of a boar in Ecuador. The finding of *L. santarosai* in testicles and semen, coupled with evidence of failed pregnancies in recipient sows is significant because it provides additional evidence of venereal transmission of leptospirosis in the swine industry. Furthermore, the silent and chronic spread *of L. santarosai* or any other species represents additional risks to public health, which needs to be approached from a one-health perspective by effective communication between animal and human health surveillance sectors.

## Supporting information

**Table S1.** Microscopic Agglutination Test of infected.

| Date | Sample | Serovar |  |  |  |  |  |
| --- | --- | --- | --- | --- | --- | --- | --- |
|  |  | Icterohaemorrhagiae | Pomona | Canicola | Hardjo | Grippotyphosa | Wolffi |
| 31/07/2019 | S1 | Negative | Negative | 1:100 | Negative | Negative | Negative |
| 19/08/2019 | S2 | Negative | 1:100 | Negative | 1:100 | Negative | Negative |

## Acknowledgments

This work was funded by the Collaboration Grants HUBI 16874 to Veronica Barragan from the Universidad San Francisco de Quito. Salary support for T. Pearson was provided by NIH/NIMHD (U54MD012388) and NIH/NIAID (R15AI156771). We are grateful to Belen Prado-Vivar (Centro de Bioinformatica, Universidad San Francisco de Quito USFQ) for her technical assistance during sequencing.

## Notes

### Competing Interest Statement

The authors have declared no competing interest.

